# Molecular basis of P[6] and P[8] major human rotavirus VP8* domain interactions with histo-blood group antigens

**DOI:** 10.1101/512301

**Authors:** Shenyuan Xu, Yang Liu, Ming Tan, Weiming Zhong, Dandan Zhao, Xi Jiang, Michael A. Kennedy

## Abstract

Initial cell attachment of rotavirus (RV) to specific cell surface glycans, which is the essential first step in RV infection, is mediated by the VP8* domain of the spike protein VP4. Recently, human histo-blood group antigens (HBGAs) have been identified as ligands or receptors for human RV strains. RV strains in the P[4] and P[8] genotypes of the P[II] genogroup share common recognition of the Lewis b and H type 1 antigens, while P[6], which is one of the other genotypes in P[II], only recognizes the H type 1 antigen. The molecular basis of receptor recognition by the major human P[8] RVs remains unknown due to lack of experimental structural information. Here, we used nuclear magnetic resonance (NMR) titration experiments and NMR-derived high ambiguity driven docking (HADDOCK) methods to elucidate the molecular basis for P[8] VP8* recognition of the Le^b^ and type 1 HBGAs and for P[6] recognition of H type 1 HBGAs. Unlike P[6] VP8* that recognizes H type 1 HGBAs in a binding surface composed of an α-helix and a β-sheet, referred as “βα binding domain”, the P[8] VP8* binds the type 1 HBGAs requiring the presence of the Lewis epitope in a previously undescribed pocket formed by two β-sheets, referred as “ββ binding domain”. The observation that P[6] and P[8] VP8* domains recognize different glycan structures at distinct binding sites supports the hypothesis that RV evolution is driven, at least in part, by selective pressure driven adaptation to HBGA structural diversity of their natural hosts living in the world. Recognition of the role that HBGAs play in driving RV evolution is essential to understanding RV diversity, host ranges, disease burden and zoonosis and to developing strategies to improve vaccines against RV infections.

**Author summary:** Rotaviruses (RV)s are the main cause of severe diarrhea in humans and animals. Significant advances in understanding RV diversity, evolution and epidemiology have been made after discovering that RVs recognize histo-blood group antigens (HBGAs) as host cell receptors. While different RV strains are known to have distinct binding preferences for HBGA receptors, the molecular basis in controlling strain-specific host ranges remains unclear. In this study, we used solution nuclear magnetic resonance to determine the molecular level details for interactions of the human P[8] and P[6] RV VP8* domains with their HBGA receptors. The distinct binding patterns observed between these major human RVs and their respective receptor ligands provides insight into the evolutionary relationships between different P[II] genotypes that ultimately determine host ranges, disease burden, zoonosis and epidemiology, which may impact future strategies for vaccine development against RVs.

## Introduction

Rotaviruses (RVs) cause severe dehydrating gastroenteritis in children younger than five years of age, accounting for two million childhood hospital admissions and leading to approximately 200,000 deaths each year [1–4]. RVs, members of the family *Reoviridae*, are double-stranded RNA viruses that contain a segmented genome, encoding six structural and six nonstructural proteins. The viral genome is encapsulated by three concentric protein layers, with the inner layer made by viral protein 2 (VP2), the intermediate shell by VP6, and the outer layer by glycosylated VP7 [5]. The protease-sensitive VP4 proteins protrude from the outer capsid shell, forming sixty spikes that are responsible for host receptor interactions. The two outer capsid proteins, VP7 and VP4, are determinants of RV G and P types, respectively, and induce G and P type-specific neutralizing antibody responses. Thus, VP7 and VP4 are used to classify RV types, which is the basis of the RV dual-nomenclature system [6]. RVs are genetically diverse, containing five genogroups (P[I]-P[V]) with 40 known genotypes (P[1]-P[40]) among group A RVs based on the VP4 sequences [7–9]. These RVs can infect humans and/or different animal species, among which P[8], P[4], and P[6] RVs in P[II] genogroup are the predominant genotypes causing diseases in humans worldwide [10–12].

RVs recognize cell surface carbohydrates through the VP4 spike proteins in a process that leads to viral attachment to the host cells in a step that is critical to initiation of host infections. Each VP4 protein is cleaved by trypsin into two domains, VP5* and VP8*, corresponding to the stalk and the distal head of the spike protein, respectively [13]. The distal VP8* head interacts with RV host glycan receptors for viral attachment, while the VP5* stalk assists in viral penetration into host cells [13,14]. Increasing data indicate that different P-type RVs exhibit distinct glycan binding specificities that may be responsible for determining host ranges as well as zoonotic transmission. While the sialidase sensitive P[1], P[2], P[3], and P[7] genotypes in the P[I] genogroup recognize sialoglycans for attachment [15–17], many sialidase insensitive RVs in the five genogroups recognize polymorphic histo-blood group antigens (HBGAs) for attachment [9,18–21]. HBGAs are complex carbohydrates that are present on red blood cells and epithelia of the gastrointestinal, respiratory and genitourinary tracts and in exocrine secretions [22,23]. HBGAs are synthesized by sequential addition of a carbohydrate moiety to a precursor disaccharide in successive steps that are genetically controlled by ABO (*ABO*, 9q34.1), H (*FUT1*, 19q13.3), secretor (*FUT2*, 19q13.3) and Lewis (*FUT3*, 19p13.3) gene families [24]. For example, the *FUT2* gene encodes an α-1,2-fucosyltransferase that catalyzes addition of an α-1,2-fucose to the precursor oligosaccharides, forming H-type antigens, while the *FUT3* gene encodes an α-1,3/4-fucosyltransferase that turns precursor oligosaccharides or H-type antigens into Le^a^ or Le^b^ antigens.

Increasing evidence supports linkages between HBGA phenotypes and RV epidemiology, indicating that HBGAs play an important role in determining interspecies transmission and disease burden of RVs. For example, whereas P[6] and P[19] RVs in the P[II] genogroup only recognize H type 1 antigen in the absence of the Lewis fucose, P[4] and P[8] RVs require the Lewis b (Le^b^) modified H type 1 antigens [19,25], supporting the observation that Le^-^ individuals may have increased risk of P[6] RV infection whereas nonsecretors and Le^-^ individuals may be resistant to P[4] and P[8] RV infections that require the Lewis and secretor fucose for host infection [26–28]. The P[9], P[14], and P[25] genotypes in the P[III] genogroup recognize type A HBGAs [9], which explains that P[14] infection in both humans and animals may due to type A HBGAs that are shared between human and animal species [9,18,29]. The P[11] RVs in the P[IV] genogroup recognize type 1 and type 2 HBGA precursors [20,21,30]. Previous studies demonstrate that bovine P[11] VP8* only recognizes type 2 HBGAs whereas human neonatal P[11] VP8* binds both type 1 and type 2 HBGAs, which is consistent with the finding that abundant type 2 glycans were found in bovine milk while both type 1 and type 2 glycans were found in human milk [30–32].

To further understand the role that HBGAs play in determining RV host ranges, evolution, and zoonosis, it is necessary to characterize the molecular basis of the interactions between VP8* domains and their glycan/HBGA receptors. The RV VP8* domain adopts a classical galectin-like fold with the central structure being an anti-parallel β-sandwich formed by a five-stranded β-sheet and a six-stranded β-sheet. Past crystallography studies have demonstrated that the VP8* domain of P[3] RVs binds sialic acids in an open-ended, shallow groove between the two β-sheets. The same binding site is used by P[7] and P[14] RVs to bind monosialodihexosylganglioside GM3 and A-type HBGAs, respectively [18,33,34]. The binding cleft is wider in P[11] than in P[3]/P[7] and P[14] strains and the binding site is shifted to span almost the entire length of the cleft between the two β-sheets. A significant advance in understanding the molecular basis of HBGA-VP8* interactions was achieved through nuclear magnetic resonance and crystallographic studies of P[19] binding to type 1 HBGA, which led to the discovery of a completely new glycan binding site composed of the N-terminal α-helix and the β-sheet containing βB, βI, βJ, βK stands [25,35]. During preparation of the manuscript, P[4] and P[6] VP8* were found to interact with H-type 1 HBGA using the same binding pocket formed by a β-sheet and the N-terminal α-helix [36].

Although the crystal structure of a P[8] human RV (Wa-like) has been reported [37], no structural data regarding the P[8] RV-glycan interactions is currently available. As an alternative to X-ray crystallography, NMR titration experiments can provide valuable information about the protein ligand interactions at the atomic level [38–40]. In this study, we employed high ambiguity driven docking (HADDOCK) using NMR chemical shift perturbation data to elucidate the molecular basis for glycan recognition by the VP8* domain of the P[8] and P[6] major human RVs. We found that P[6] VP8* bound the tetra-(LNT) and penta-(LNFP I) saccharide of type 1 HBGAs without Lewis epitopes in the open pocket formed by the α-helix and the β-sheet consisting of βK, βJ, βI, βB strands. However, P[8] VP8* bound the Lewis b (Le^b^) tetra-saccharide and a Le^b^-containing hexa-saccharide (LNDFH I), in the shallow cleft formed by the two twisted antiparallel β-sheets. The discovery and documentation of the complex and variable receptor binding patterns of the four P[II] RVs (P[4], P[6], P[8[and P[19]) reported here and elsewhere continues to expand our understanding of the evolution, host ranges, disease burden and zoonosis of P[II] RVs.

## Results

### Identification of specific HBGA glycan interactions with P[8] VP8*

P[8] VP8* has been previously reported to bind the Le^b^ tetra-saccharide and the hexa-saccharide LNDFH I that contains the Lewis epitope (Fig 1a) [25], however structures of their complexes do not exist at this time. Here, proton saturation transfer NMR experiments were used to characterize the specific interactions between the P[8] RV VP8* domain and the Le^b^ tetra-saccharide and hexa-saccharide LNDFH I HBGAs. Proton NMR resonances unique to the protein were selectively saturated and saturation transfer to protons on the bound ligand measured using STD NMR experiments. Protons of glycan sugar moieties that displayed efficient saturation transfer, i.e. large STD amplification factors, were inferred to be in direct contact with the VP8* surface, whereas protons of glycan sugar moieties free from saturation transfer were assumed to be pointing away from the protein surface [41,42]. Complete glycan proton chemical shift assignments enabled determination of HBGA saccharide moieties in direct contact with the protein (Fig S1). P[8] VP8* was found to bind both the Le^b^ tetra-saccharide (Fucα1-2Galβ1-3[Fucα1-4]GlcNAc) and LNDFH I (Fucα1-2Galβ1-3[Fucα1-4]GlcNAcβ1-3Galβ1-4Glc) (Fig 2). Protons in the type 1 precursor Galβ1-3GlcNAc motif of both the Le^b^ tetra-saccharide (Fig 2b) and LNDFH I (Fig 2d) experienced strong magnetization saturation transfer, indicating that the type 1 precursor Galβ1-3GlcNAc motif was important for P[8] VP8* recognition in both HBGA glycans. The secretor fucose and Lewis fucose moieties of Le^b^ tetra-saccharide and LNDFH I (Fig 1) were also involved in binding interactions, however, the secretor fucose in both Le^b^ tetra-saccharide and LNDFH I, and Lewis fucose in LNDFH I received relatively weak saturation transfer, while the Lewis epitope in Le^b^ tetra-saccharide experienced relatively high magnetization transfer. The galactose (Gal-IV) and glucose (Glc-V) moieties of LNDFH I did not appear to make direct or strong contact with the P8 VP8* surface as their protons experienced relatively weak saturation transfer. Analysis of the NMR titration experiments indicated that P[8] VP8* bound more tightly to LNDFH I with a dissociation constant (Kd) of 6.9 mM compared to Le^b^ tetra-saccharide, which had a much higher Kd value of 43.4 mM (Fig S4).

**Fig 1.**
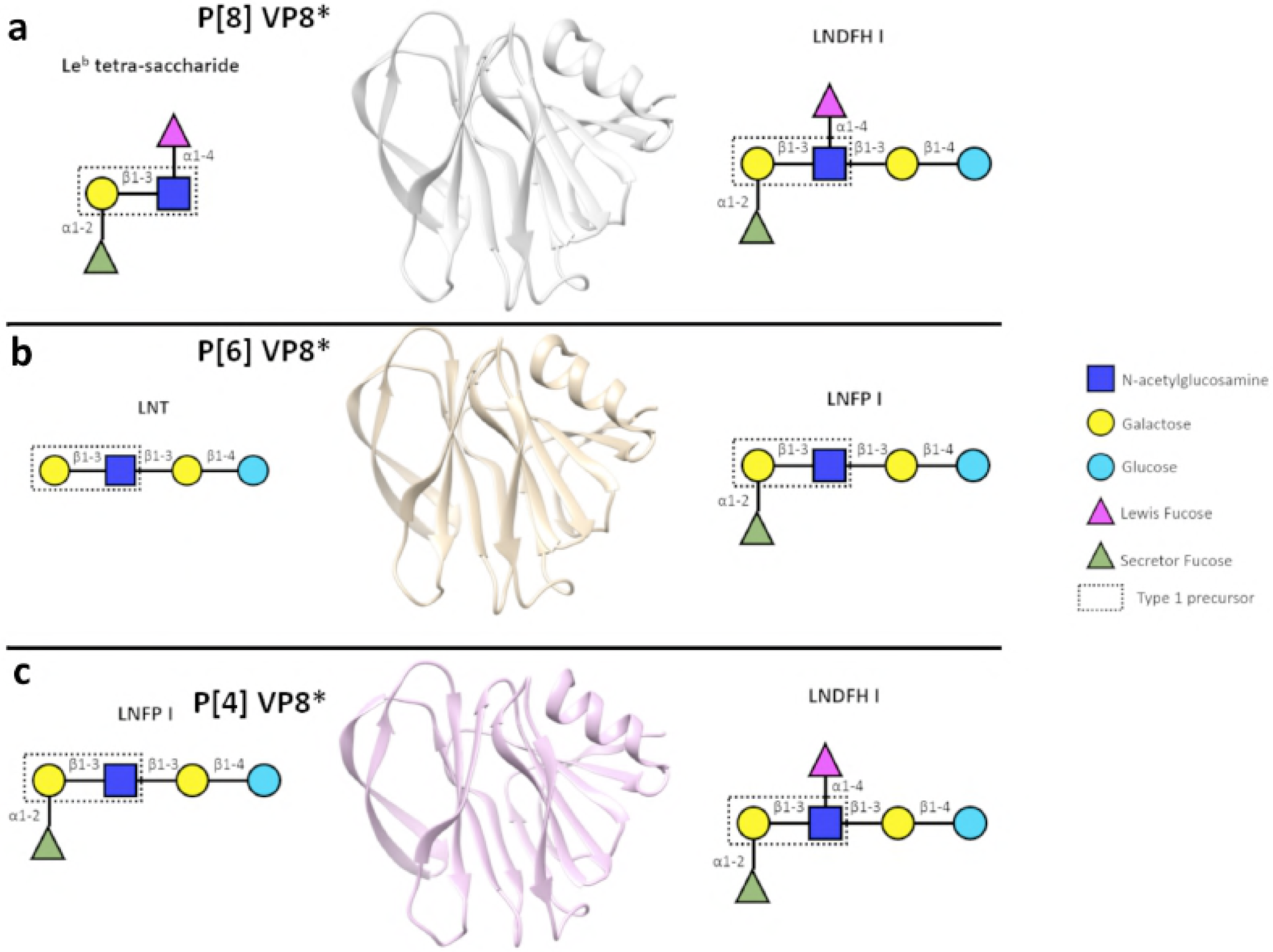
Schematic representation of glycan ligands used for NMR titration experiments. (a) Schematic representation of P[8] VP8* and its binding glycan structures, Le^b^ tetra-saccharide and LNDFH I. (b) Schematic representation of P[6] VP8* and its binding glycan structures, LNT and LNFP I. (c) Schematic representation of P[4] VP8* and its binding glycan structures, LNDFH I and LNFP I.

**Fig 2.**
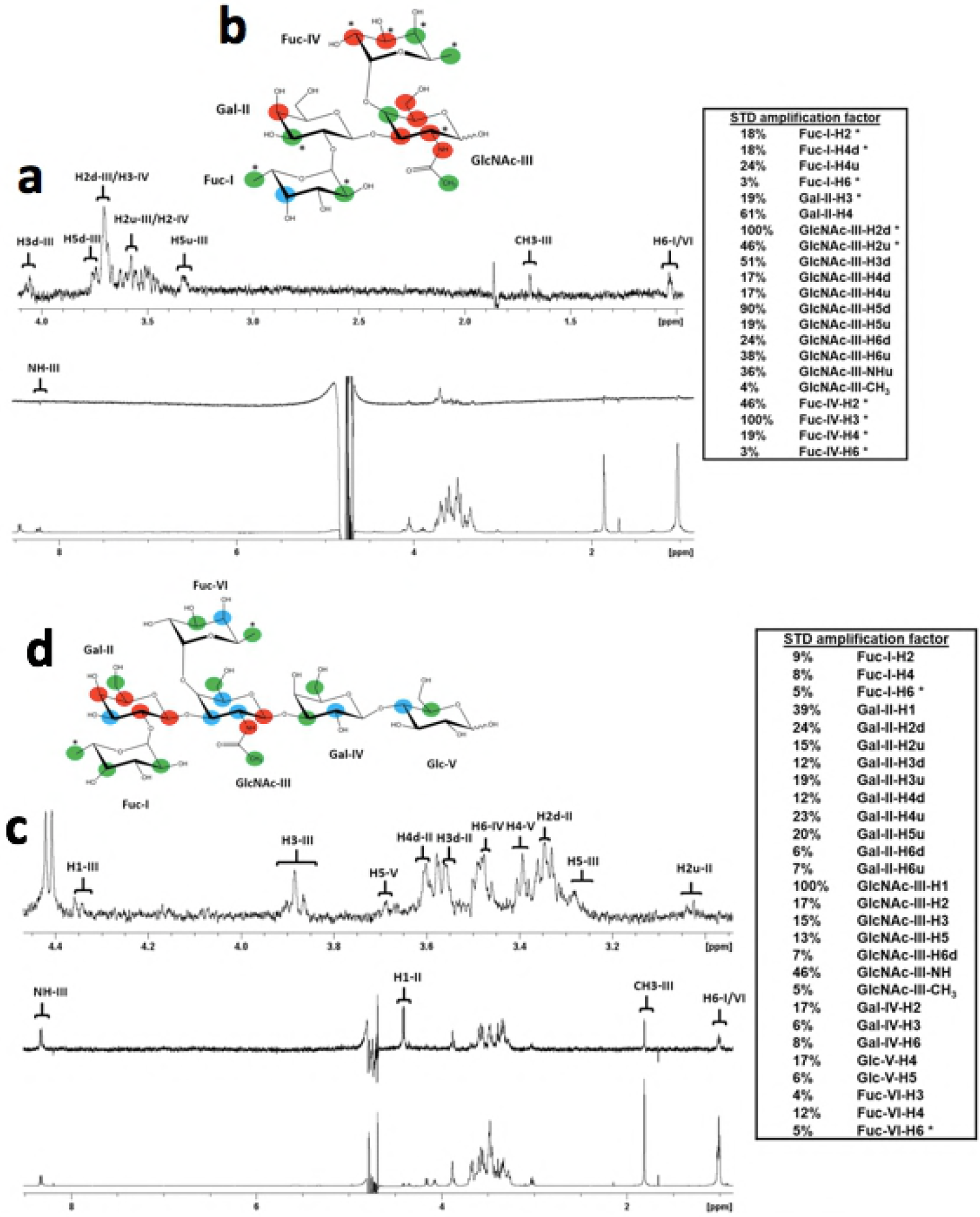
STD NMR analysis of Le^b^ tetrasaccharide and LNDFH I glycan interactions with the P[6] VP8* domain. (a) NMR spectra of Le^b^ tetra-saccharide in complex with P[8] VP8*. The bottom spectrum in (a) is ^1^H NMR reference spectrum of P[8] VP8* (33 μM) with Le^b^ tetra-saccharide (1.86 mM). The middle spectrum in (a) is STD NMR spectrum of P[8] VP8* (33 μM) with Le^b^ tetra-saccharide (1.86 mM). The protein was saturated with a cascade of 40 Gaussian-shaped pulses at 0.11 p.p.m., and the off-resonance was set to 50 ppm. The upper spectrum in (a) is the expansion of the STD NMR from 4.1 to 0.9 ppm. (b) The epitope mapping of Le^b^ tetra-saccharide when bound to P[8] based on the STD effects: red, strong STD NMR effects (>30%); blue, medium STD NMR effects (20%-30%); green, weak STD NMR effects (<20%). (c) NMR spectra pf LNDFH I in complex with P[8] VP8*. The bottom spectrum in (c) is ^1^H NMR reference spectrum of P[8] VP8* (46 μM) with LNDFHI (2.3 mM). The middle spectrum in (c) is STD NMR spectrum of P[8] VP8* (46 μM) with LNDFHI (2.3 mM). The protein was saturated with a cascade of 40 Gaussian-shaped pulses at -0.25 ppm, and the off-resonance was set to 50 ppm. The upper spectrum in (c) is the expansion of the STD NMR from 4.5 to 3.0 p.p.m. (d) The epitope mapping of LNDFH I when bound to P[8] based on the STD effects: red, strong STD NMR effects (>20%); blue, medium STD NMR effects (10%-20%); green, weak STD NMR effects (<10%). “*” means the overlapping of signals in the STD NMR spectrum. “d” indicates a downfield proton, “u” indicates an upfield proton.

### Identification of specific HBGA glycan interactions with P[6] VP8*

P[6] VP8* has been previously reported to bind LNT and LNFP I (Fig 1b) [25], and the crystal structure of the P[6] VP8* domain complex with LNFP I has been reported (ref), however its structure with LNT has not been reported at this time. Here, proton saturation transfer NMR experiments were used to characterize the interactions between the P[6] RV VP8* domain and the LNT and LNFP I HBGAs. STD NMR experiments indicated that P[6] VP8* recognized both LNT (Galβ1-3GlcNAcβ1-3Galβ1-4Glc) (Fig 1b) and LNFP I (Fucα1-2Galβ1-3GlcNAcβ1-3Galβ1-4Glc) (Fig 1b) glycans (Fig 3). Protons in the Galβ1-3GlcNAcβ1-3Gal chain of both LNT and LNFP I experienced strong saturation transfer indicating that this motif, common to both LNT and LNFP 1, was important in P[6] VP8*-HBGA interactions. The terminal glucose of LNT and LNFP I also experienced saturation transfer from the protein, although the transfer was unevenly distributed in the glucose between the two glycans, with H1 in the Glc-V moiety of LNFP I having stronger saturation transfer. Fuc-I in LNFP I appeared to point away from the protein surface as its protons received only weak saturation transfer. The affinity of P[6] VP8* to LNT was higher (Kd = 4.3 mM) compared to that of LNFP I (Kd = 28.2 mM), based on the analysis of the NMR titration data (Fig S6).

**Fig 3.**
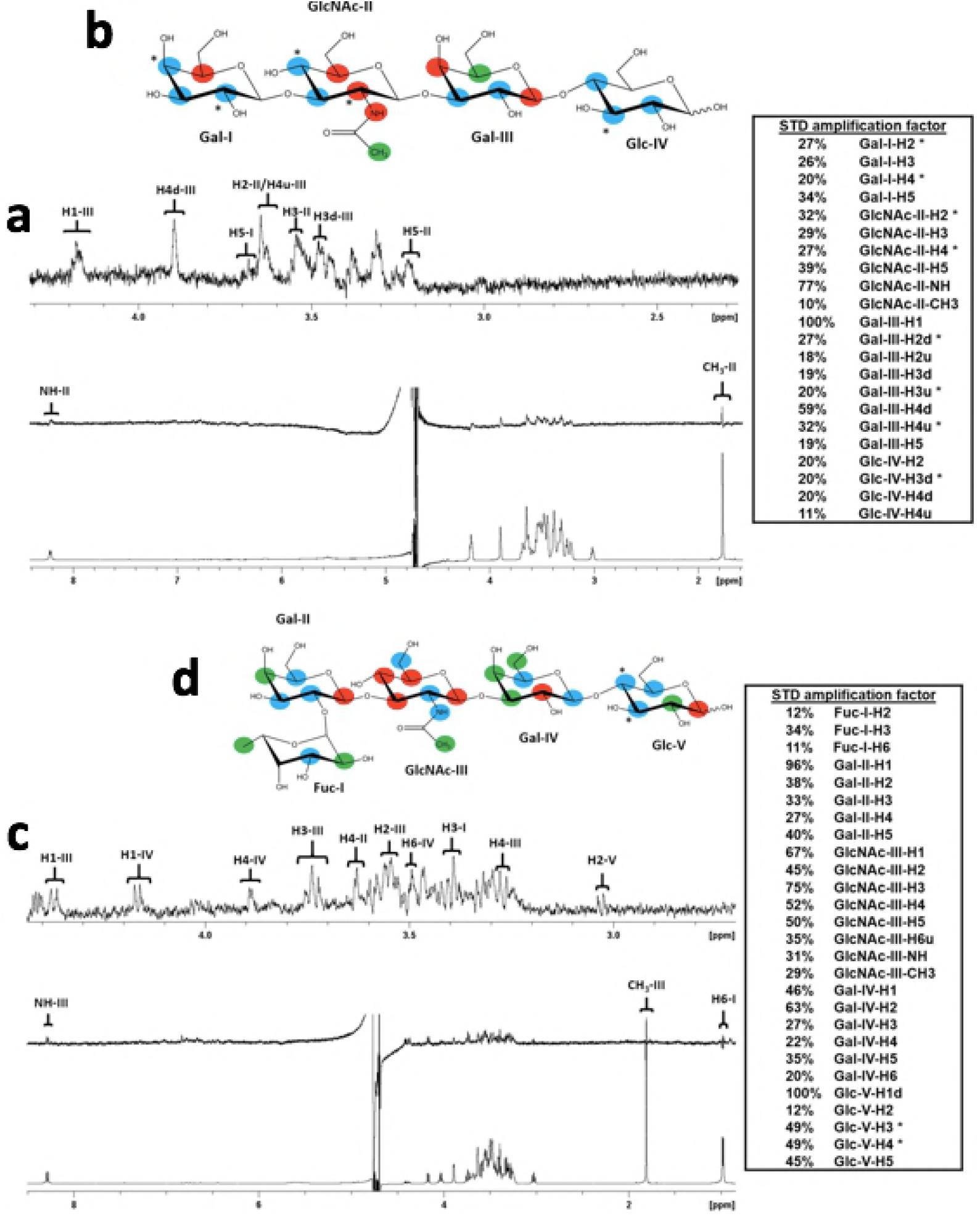
STD NMR analysis of LNT and LNFPI glycan interactions with the P[6] VP8* domain. (a) NMR spectra pf LNT in complex with P[6] VP8*. The bottom spectrum in (a) is the ^1^H NMR reference spectrum of P[6] VP8* (46 μM) with LNT (4.6 mM). The middle spectrum in (a) is the STD NMR spectrum of P[6] VP8* (46 μM) with LNT (4.6 mM). The protein was saturated with a cascade of 40 Gaussian-shaped pulses at 0.63 ppm, and the off-resonance was set to 50 ppm. The upper spectrum in (a) is the expansion of the STD NMR from 4.3 to 2.3 ppm. (b) The epitope map of LNT when bound to P[6] based on the STD effects: red, strong STD NMR effects (>30%); blue, medium STD NMR effects (20%-30%); green, weak STD NMR effects (<20%). (c) NMR spectra pf LNFPI in complex with P[6] VP8*. The bottom spectrum in (c) is the ^1^H NMR reference spectrum of P[6] VP8* (46 μM) with LNFPI (2.3 mM). The middle spectrum in (c) is the STD NMR spectrum of P[6] VP8* (46 μM) with LNFPI (2.3 mM). The protein was saturated with a cascade of 40 Gaussian-shaped pulses at -0.21 ppm, and the off-resonance was set to 50 ppm. The upper spectrum is the expansion of the STD NMR from 4.4 to 1.0 ppm. (d) The epitope map of LNFPI when bound to P[6] based on the STD effects: red, strong STD NMR effects (>50%); blue, medium STD NMR effects (30%-50%); green, weak STD NMR effects (<30%). “*” means the overlap of signals in the STD NMR spectrum. “d” indicates a downfield proton, “u” indicates an upfield proton.

### Mapping HBGA glycan interactions onto the surface of P[8] VP8* using chemical shift perturbations observed in HSQC NMR titration experiments

In order to identify the HBGA glycan binding surface on the P[8] VP8* domain, the P[8] VP8* backbone resonances were assigned using triple resonance NMR experiments and the backbone chemical shifts were deposited to the BioMagResBank database (accession number 27598). The 2D ^l·^H-^15^N HSQC spectrum of P[8] VP8* was of excellent quality (Fig S2) and the backbone assignments for NH pairs were 94% complete. Titrations of Le^b^ tetra-saccharide and LNDFH I to ^15^N-labeled P[8] VP8* caused significant chemical shift perturbations and disappearance of some resonances, while the chemical shifts of the majority signals were slightly or not affected (Fig 4a,d). Complete profiles of chemical shift perturbations of P[8] VP8* upon addition of Le^b^ tetra-saccharide and LNDFH I are shown in (Fig S3). Titration with either Le^b^ tetra-saccharide or LNDFH I resulted in a common pattern of affected residues, with L157 of βH strand, K168 of βI strand, G178, E179, and A183 of βJ-K loop, and D186 of βK strand exhibiting large chemical shifts changes (Fig 4). In addition to the commonly affected residues, the two glycans also exhibited some slightly different chemical shift perturbations. For example, addition of the Le^b^ tetra-saccharide, caused large chemical shifts changes clustered in the common area, including T156 of βH strand and F176 of βJ strand, and A107 of βD strand, however, G145 of βG-H loop that was distant from the common area, also showed large chemical shifts changes (Fig 4b,c). Upon addition of LNDFH I, some amino acids with significant chemical shifts perturbations that clustered in the common protein surface included R154 of βH strand, H177 of βJ strand, T185 of βK strand, plus several residues that were not clustered with the other group of residues experienced large chemical shifts perturbations, including T78 and D79 of βA-B loop, T161 of βH-I loop, V164 and G165 of βI strand, I207 of βM strand (Fig 4e, f).

**Fig 4.**
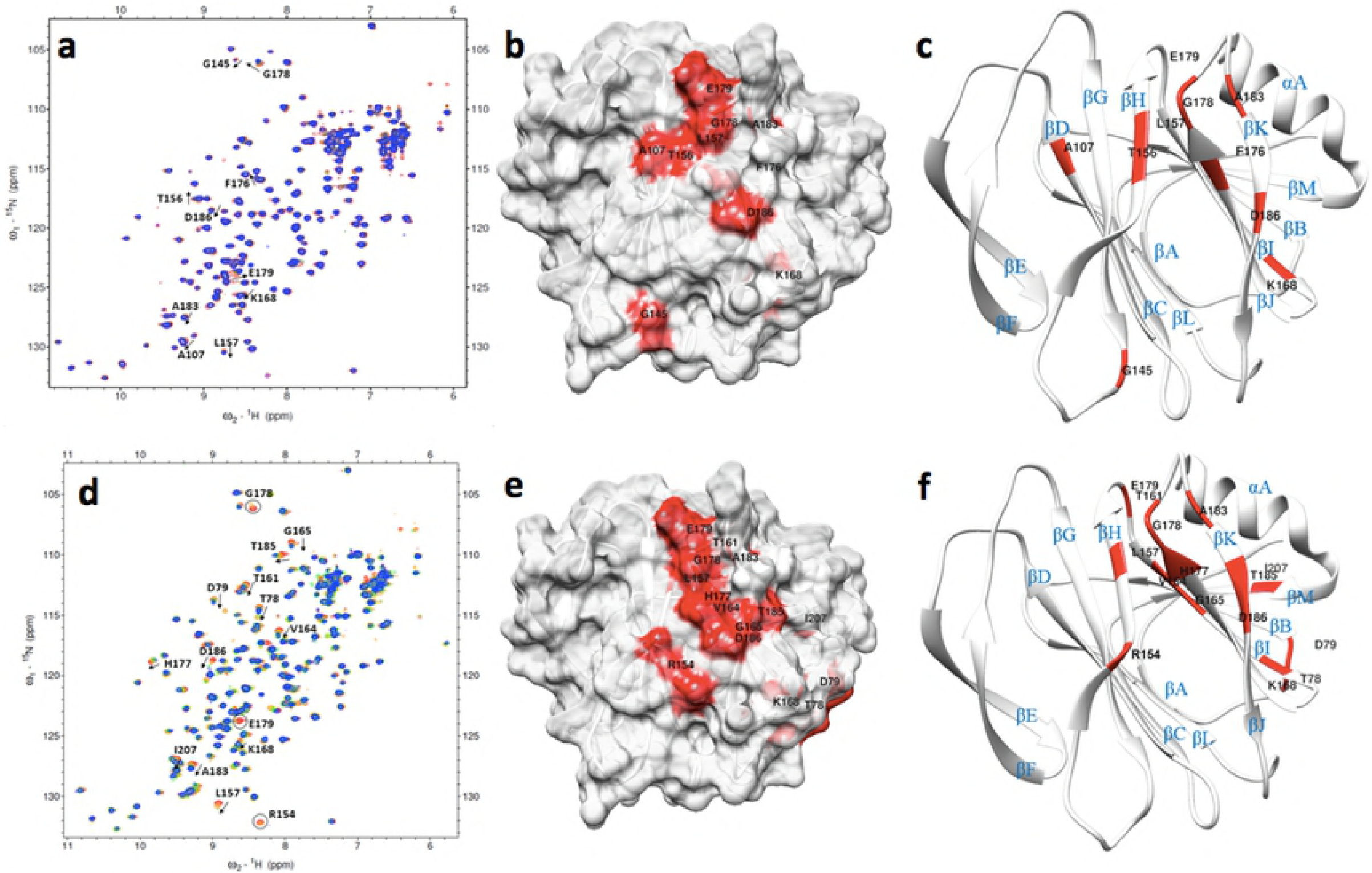
NMR titration analysis of P[8] VP8* domain with Leb and LNDFH I glycans. (a) Chemical shift changes in P[8] VP8* [PDB ID: 2DWR] upon addition of Le^b^ tetra-saccharide. Titrations were followed by acquisition of two-dimensional ^1^H-^15^N HSQC spectra of 0.31 mM ^15^N-labeled P[8] VP8* collected at 283 K. The NMR data correspond to increasing ligand/protein ratios of 0:1 (red), 8:1 (orange), 18:1 (green), and 30:1 (blue). (b) Location of the large chemical shift changes on the P[8] VP8* surface [PDB ID: 2DWR] upon binding to Le^b^ tetra-saccharide. Amino acids with chemical shift changes greater than 2.5σ or disappeared after titration are colored with red. (c) Ribbon Diagram shows the location of large chemical shift changes on the P[8] VP8* surface upon binding to Le^b^ tetra-saccharide. The secondary structure is labeled. (d) Chemical shift changes in P[8] VP8* upon addition of LNDFH I. Titrations were followed by acquisition of two-dimensional ^1^H-^15^N HSQC spectra of 0.20 mM ^15^N-labeled P[8] VP8* collected at 298 K. The NMR data correspond to increasing ligand/protein ratios of 0:1 (red), 6:1 (orange), 18:1 (green), and 25:1 (blue). The arrows show residues that have chemical shifts changes upon titration. Peaks that disappeared upon titration are circled. (e): Location of the large chemical shift changes on the P[8] VP8* surface [PDB ID: 2DWR] upon binding to LNDFH I. Amino acids with chemical shift changes greater than 1.5σ or disappeared after titration are colored with red. (f): Ribbon Diagram shows the location of large chemical shift changes on the P[8] VP8* surface upon binding to LNDFH I. The secondary structure is labeled.

### Mapping HBGA glycan interactions onto the surface of P[6] VP8* using chemical shift perturbations observed in HSQC NMR titration experiments

The P[6] VP8* backbone resonances were assigned using triple resonance experiments and the backbone chemical shifts were deposited to the BioMagResBank database (accession number 27591). The 2D ^1^H-^15^N HSQC spectrum of the P[6] VP8* domain was excellent (Fig S2) and the backbone assignments for NH pairs were 92% complete. Titration of ^15^N-labeled P[6] VP8* with LNT (Fig 1b) and LNFP I (Fig 1b) caused some resonances to experience significant chemical shift perturbations or to disappear altogether, while the remaining peaks were weakly or not affected (Fig 5, Fig S5). Addition of LNT caused significant chemical shifts perturbations clustered in the region composed of the following residues: N78 of the βA-B loop, Y80 of βB strand, F169 of βI strand, W174 of βJ strand, T184 and T185 of βK strand, Q211, E212, K214, C215, S216 of αA helix (Fig 5b,c). In addition, E141 of βG strand, T190, S191, and N192 of βK-L loop, Q89 of βB-C loop, H201 of βL-M loop, which were not in the main clustered region, also showed large chemical shifts changes. Upon adding LNFP I (Fig 1b), residues that showed significant changes of chemical shifts were clustered in the area that consisted of N78 of βA-B loop, Y80 of βB strand, F169 of βI strand, V173 and W174 of βJ strand, T184, T185, D186, and S188 of βK strand, and Q211, E212, K214, C215, S216 of αA helix (Fig 5e,f). In addition, T190, S191, N192, and L193 of βK-L loop had large chemical shifts perturbation upon adding LNFP I.

**Fig 5.**
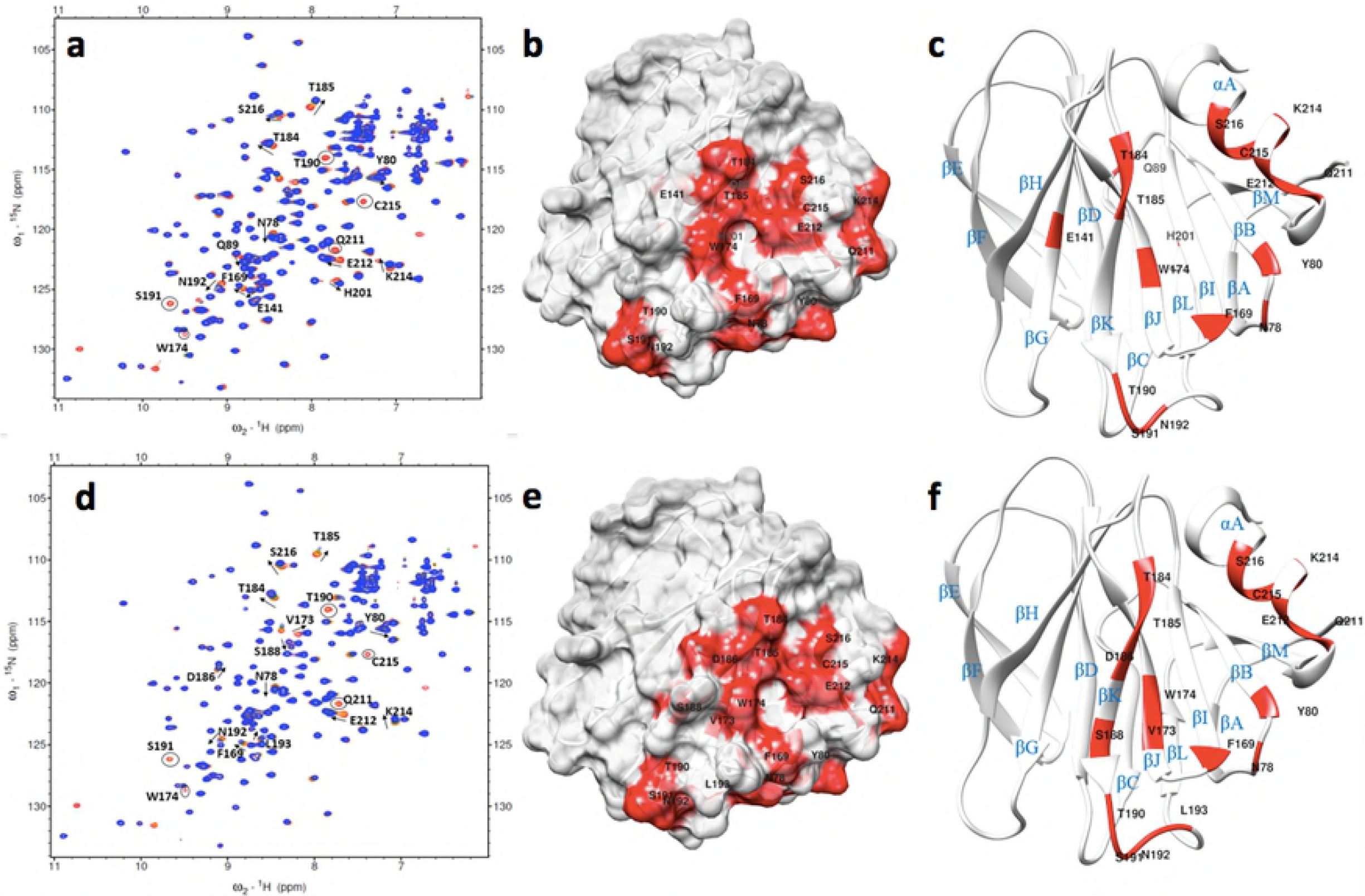
NMR titration analysis of P[6] VP8* domain with LNT and LNFP I glycans. (a) Chemical shift changes in P[6] VP8* [PDB ID: 5YMU] upon addition of LNT. Titrations were followed by acquisition of two-dimensional ^1^H-^15^N HSQC spectra of 0.334 mM^15^N-labeled P[6] VP8* collected at 283 K. The NMR data correspond to increasing ligand/protein ratios of 0:1 (red), 8:1 (orange), 18:1 (green), and 25:1 (blue). (b) Location of the large chemical shift changes on the P[6] VP8* surface [PDB ID: 5YMU] upon binding to LNT. Amino acids with chemical shift changes greater than 2σ or disappeared after titration are colored with red. (c) Ribbon Diagram shows the location of large chemical shift changes on the P[8] VP8* surface upon binding to Le^b^ tetra-saccharide. The secondary structure is labeled. (d) Chemical shift changes in P[6] VP8* upon addition of LNFP I. Titrations were followed by acquisition of two-dimensional ^1^H-^15^N HSQC spectra of 0.241 mM ^15^N-labeled P[6] VP8* collected at 283 K. The NMR data correspond to increasing ligand/protein ratios of 0:1 (red), 8:1 (orange), 18:1 (green), and 35:1 (blue). The arrows show residues that have chemical shifts changes upon titration. Peaks that disappeared upon titration are circled. (e) Location of the large chemical shift changes on the P[6] VP8* surface [PDB ID: 5YMU] upon binding to LNFPI. Amino acids with chemical shift changes greater than 1.5σ or disappeared are colored with red. (f) Ribbon Diagram shows the location of large chemical shift changes on the P[8] VP8* surface upon binding to LNDFH I. The secondary structure is labeled.

### Models of the structures of the P[8] VP8* domain bound to the glycan receptor ligands

Superposition of the top five best-scoring P[8] VP8* structures from HADDOCK docking showed that the bound structures of the Le^b^ tetra-saccharide and LNDFH I clustered in the same shallow cleft formed by the edges of two β-sheets (referred to hereafter as the “ββ binding domain”) with the only exception being that LNDFH I covered more surface area (Fig 6 & Fig S7). The ββ binding domain was mainly composed of the βH strand, the C-terminus part of βJ strand, the N-terminus part of βK strand, and the βJ-K loop connecting the two strands. The models of P[8]-Le^b^ and P[8]-LNDFH I with the best HADDOCK scores were chosen for detailed analysis (Fig 6& FigS8). In the P[8]-Le^b^ tetra-saccharide complex, the secretor epitope Fuc-I (Fig 1a) pointed away from the protein surface (Fig 6a), Gal-II appeared to form hydrogen bonds with E179, R182, and T180 and hydrophobic interactions with P181, GlcNAc-III formed hydrophobic interactions with H177, G178, A183 and T184 and a hydrogen bond with R182, and the Lewis epitope Fuc-IV inserted into the hydrophobic core formed by K138, T156, L157, and T158. The orientation of the type 1 precursor Galβ1-3GlcNAc (Fig 1) was different in the P[8]-LNDFH I complex compared to in the Le^b^-bound P[8] structure, such that the Lewis epitope pointed away from the protein surface (Fig 6d). Regarding the other residues, the secretor epitope Fuc-I had hydrophobic interactions with K138, T158, E179 and formed hydrogen bonds with R182, Gal-II formed hydrophobic interactions with A183 and a hydrogen bond with T184, GlcNAc-III contacted H177 and G178 through hydrophobic interactions, Gal-IV formed hydrophobic interactions with R155 and T156 and a hydrogen bond with N153, and Glc-V contacted Y152 via hydrophobic interactions and was further stabilized by hydrogen bonds with N153 and R154.

**Fig 6.**
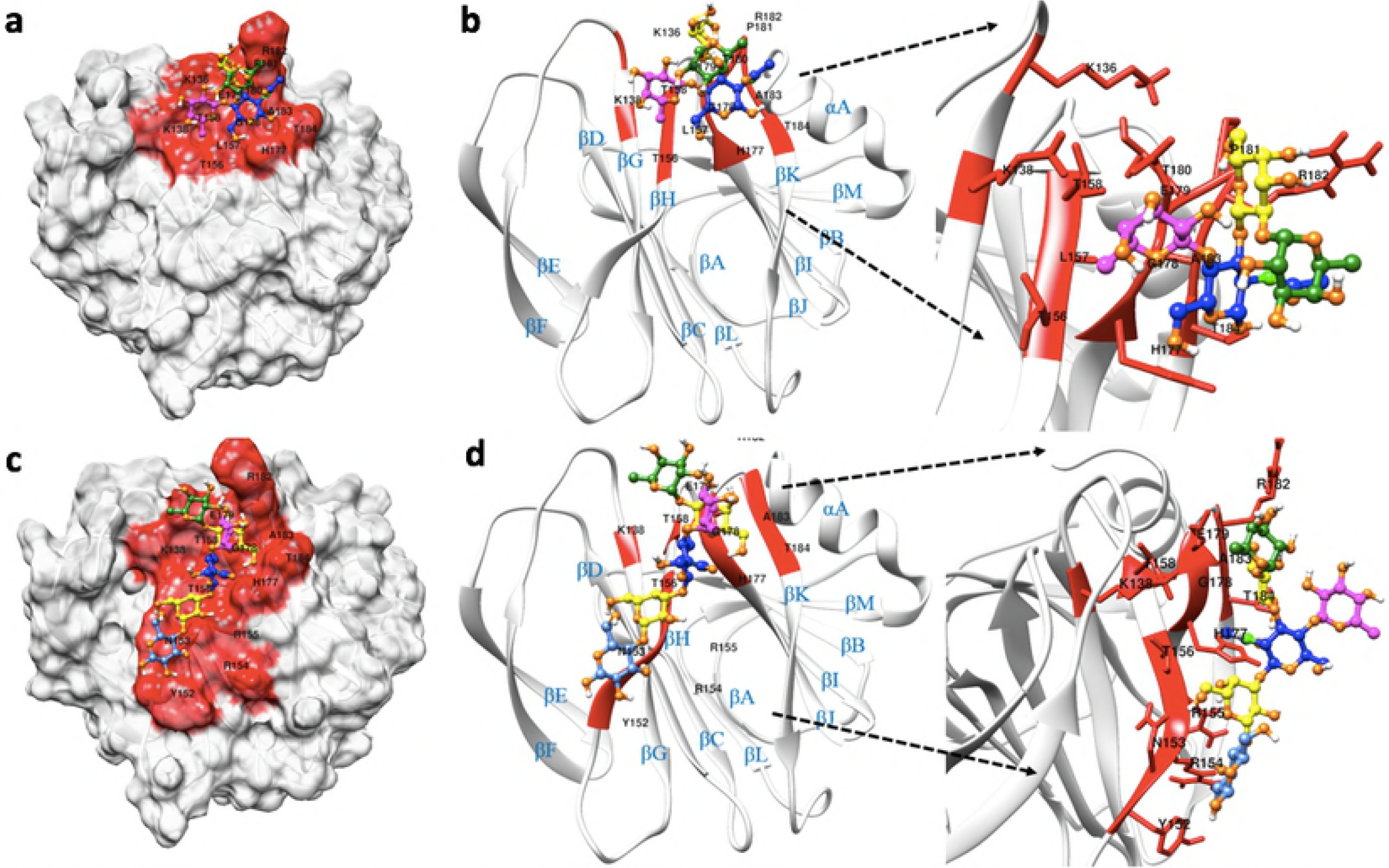
NMR-driven HADDOCK models of the P[8] VP8* domain bound to Le^b^ and LNDFH I. (a) The HADDOCK docking result for the interaction of P[8] VP8* and Le^b^ tetra-saccharide. The cartoon shows the complex conformation with the best HADDOCK score. (b) Ribbon Diagram shows the Le^b^ tetra-saccharide-bound P[8] VP8* with the best HADDOCK score. The right panel in (b) highlights the binding pocket of P[8] VP8* in recognizing Le^b^ tetra-saccharide. (c) The HADDOCK docking result for the interaction of P[8] VP8* and LNDFH I. The cartoon shows the lowest-energy conformation with the best HADDOCK score. (d) Ribbon Diagram shows the LNDFH I-bound P[8] VP8* with the best HADDOCK score. The right panel in (d) highlights the binding pocket of P[8] VP8* in recognizing LNDFH I. Red represents the binding interface. Different colors were used to represent different carbohydrate moiety of the ligand: magenta colors the Lewis Fucose residue, green colors the secretor Fucose residue, yellow colors the Galactose residue, blue colors the N-Acetylglucosamine residue, cornflower blue colors the Glucose residue.

### Models of the structures of the P[6] VP8* domain bound to the glycan receptor ligands

The top five best-scoring P[6] VP8* structures revealed that LNT and LNFP I clustered in the same shallow groove composed of the N-terminal α-helix and the β-sheet composed of the βK, βJ, βI, βB strands (hereafter referred to as the “βα binding domain”) (Fig S9). The binding interface generated by HADDOCK docking was in agreement with the binding region revealed by the X-ray crystal structure of LNFP I-bound P[6] (strain RV3) (Fig S9c). One noticeable difference was the nearly 180 degrees rotation about the glycosidic bond of GlcNAcβ1-3Gal, leading to the terminal Glc epitope being opposite in the NMR docking structure compared to crystal structure (Fig S9d). The best scoring model in the best cluster was chosen for further analysis (Fig 7, Fig S10). The Galβ1-3GlcNAc type 1 precursor in both the LNT-bound and LNFP I-bound P[6] VP8* inserted into the pocket formed by W174, T184, T185, R209, E212, S213, and S216, and appears to be stabilized through hydrogen bonding and hydrophobic interactions. In the LNT-bound P[6] VP8*, the Gal-III contacted F169, Y170, W174 through hydrophobic interactions, and the Glc-IV pointed away from the protein surface. In the LNFP I-bound P[6] VP8*, the Fuc-I was stabilized by R209 and S210 through hydrophobic interactions and the terminal Galβ1-4Glc epitope was stabilized by the side chains of M167, F169, Y170, W174.

**Fig 7.**
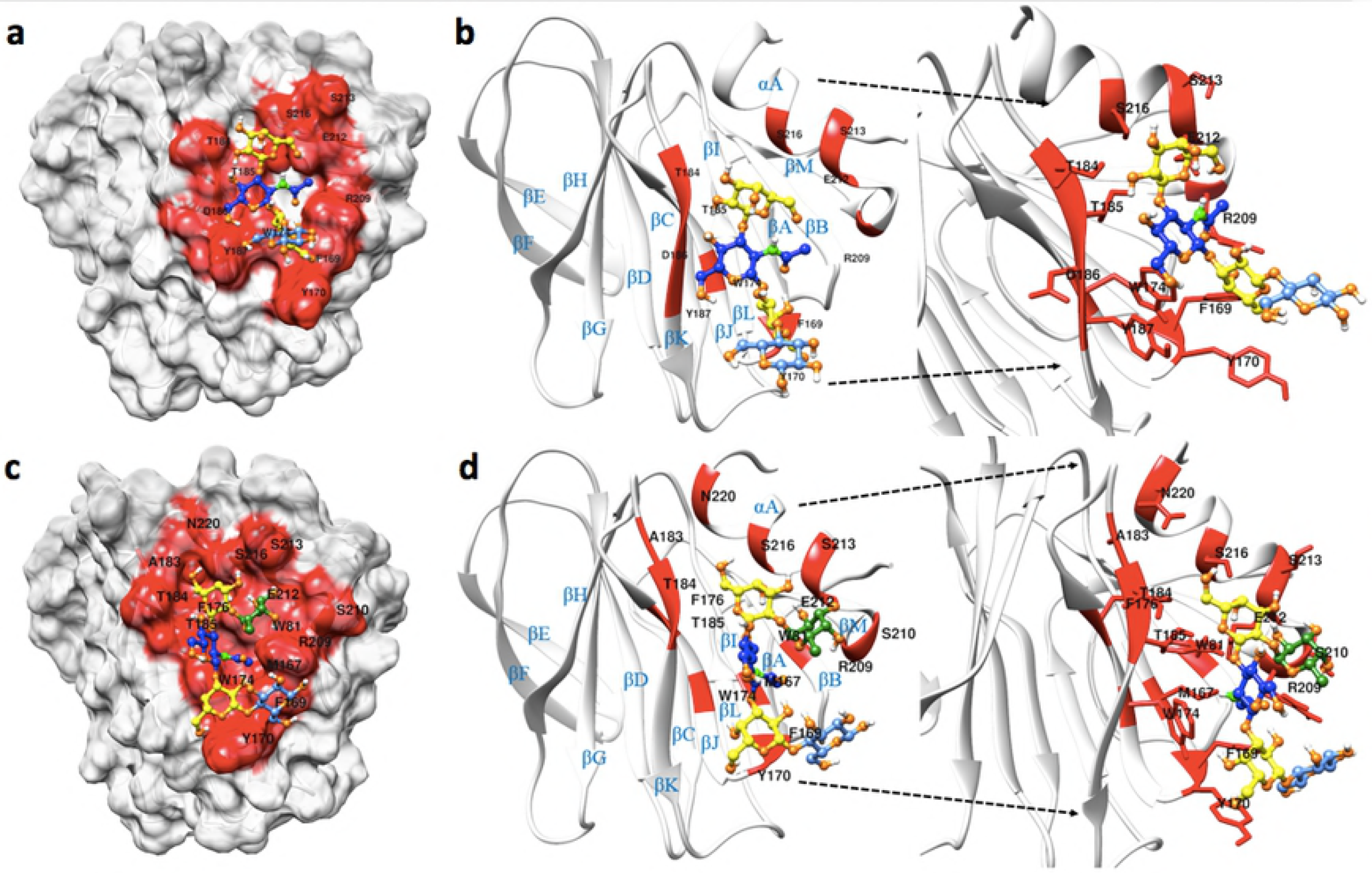
NMR-driven HADDOCK models of the P[6] VP8* domain bound to LNT and LNFP I. (a) The HADDOCK docking result for the interaction of P[6] VP8* and LNT. The cartoon shows the model with the best HADDOCK in the best cluster. (b) Ribbon Diagram shows the LNT-P[6] VP8* with the best HADDOCK score. The right panel in (b) highlights the binding pocket of P[6] VP8* in recognizing LNT. (c) The HADDOCK docking result for the interaction of P[6] VP8* and LNFP I. The cartoon shows the model with the best HADDOCK in the best cluster. (d) Ribbon Diagram shows the LNFP I-P[6] VP8* with the best HADDOCK score. The right panel in (d) highlights the binding pocket of P[6] VP8* in recognizing LNFP I. Red represents the binding interface. Different colors were used to represent different carbohydrate moiety of the ligand: Green colors the secretor Fucose residue, yellow colors the Galactose residue, blue colors the N-Acetylglucosamine residue, cornflower blue colors the Glucose residue.

### Structural conservation and variations within the VP8* glycan binding pocket

When the sequences of different genotypes of VP8* with known structures and experimentally documented binding interfaces were aligned (Fig 8), it was found that RV VP8*s use one of two common binding pockets to accommodate different glycans (Fig 9), either the ββ binding domain or the βα binding domain, however, slight sequence and structural variations among different genotypes cause the glycans to bind differently within the binding pockets in a genotype-specific manner (Fig 9). P[3]/P[7] in P[I] and P[14] RVs in P[III] use the ββ binding domain as a glycan binding interface in which R101 in the βC-D loop and motif 187-XYYX-190 in the βK strand are highly conserved (Fig 8). The conserved pattern observed in P[I] and P[III] genogroups was not found in other genogroups. For example, compared to R101 and Y188 in P[3]/P[7] and P[14], amino acids 101 and 188 are phenylalanine and threonine respectively in the P[11] VP8* in the P[IV] genogroup, whose binding interface covers almost the entire length of βH strand and the C-terminal of βJ strand as well as the βJ-K loop (Fig 9c). For P[4], P[6] and P[19] in the P[II] genogroup, the binding interface is shifted to the βα binding domain (Fig 8, 9). Some key residues involved in the binding pocket in the P[II] genogroup are highly conserved, e.g. W81 in the βB strand, W174 in the βJ strand, T184 and T185 in the βK strand, R209 in the βm-αA loop, and E212 in the αA helix (Fig 8, 9). While P[4] and P[8] RVs are genetically closely related, P[8] employed the ββ binding domain to bind the Le^b^ tetra-saccharide and LNDFH I whereas P[4] used the βα binding domain to recognize LNFP I that contained no Lewis epitope (Fig 9d,e). It appears that the addition of Lewis epitope may cause P[8] to shift LNDFH I binding to the ββ binding domain. P[8] showed no binding to LNFP I in our STD experiment, which contains no Lewis epitope, indicating that the Lewis epitope is necessary for P[8] RVs to recognize HBGAs (Fig S11). Also based on our STD results, P[4] could recognize both LNFP I that has no Lewis epitope and LNDFH I that contains the Lewis epitope (Fig S12). We therefore hypothesized that P[4] will bind the Lewis-containing LNDFH I in the ββ binding domain, just as P[8] did, and we have indicated this possibility in Fig 9. It was also deduced that Wa strain of P[8] can recognize the Le^b^ and H type 1 glycans in the same pattern as the BM13851 strain of P[8] since the amino acids involved in the binding pocket are same.

**Fig 8.**
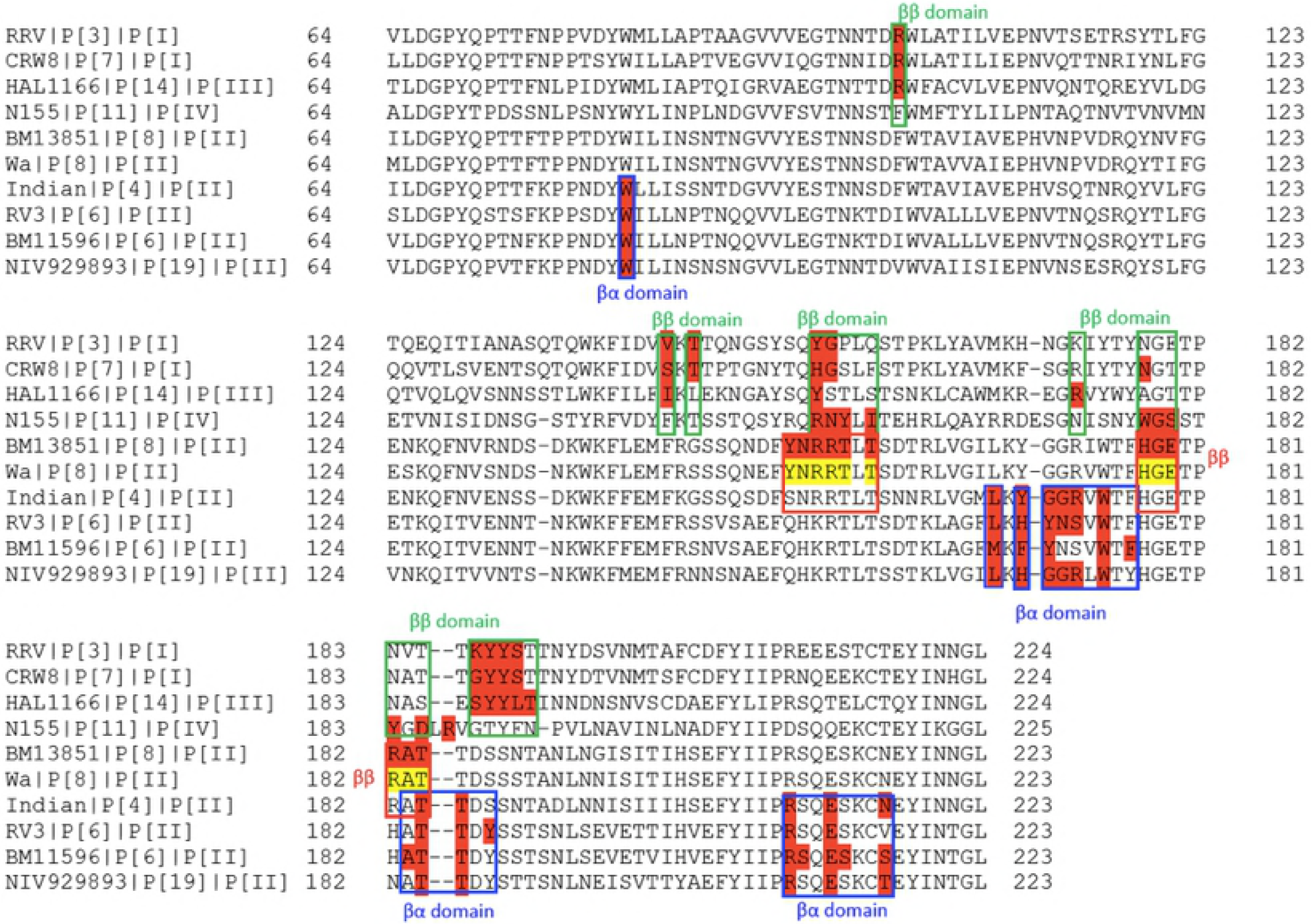
Structure-based sequence alignment of VP8* domains. The VP8* sequences of RVs with known binding sites are included. The VP8* sequences of P[19], P[6], P[4] and P[8] in P[II] genogroup, P[3], P[7] in P[I] genogroup, P[14] in P[III] genogroup and P[11] in P[IV] genogroup are included and the amino acids corresponding to the binding site are highlighted in red. Yellow highlights the potential binding site of Wa strain of P[8] VP8* in recognizing HBGAs.

**Fig 9.**
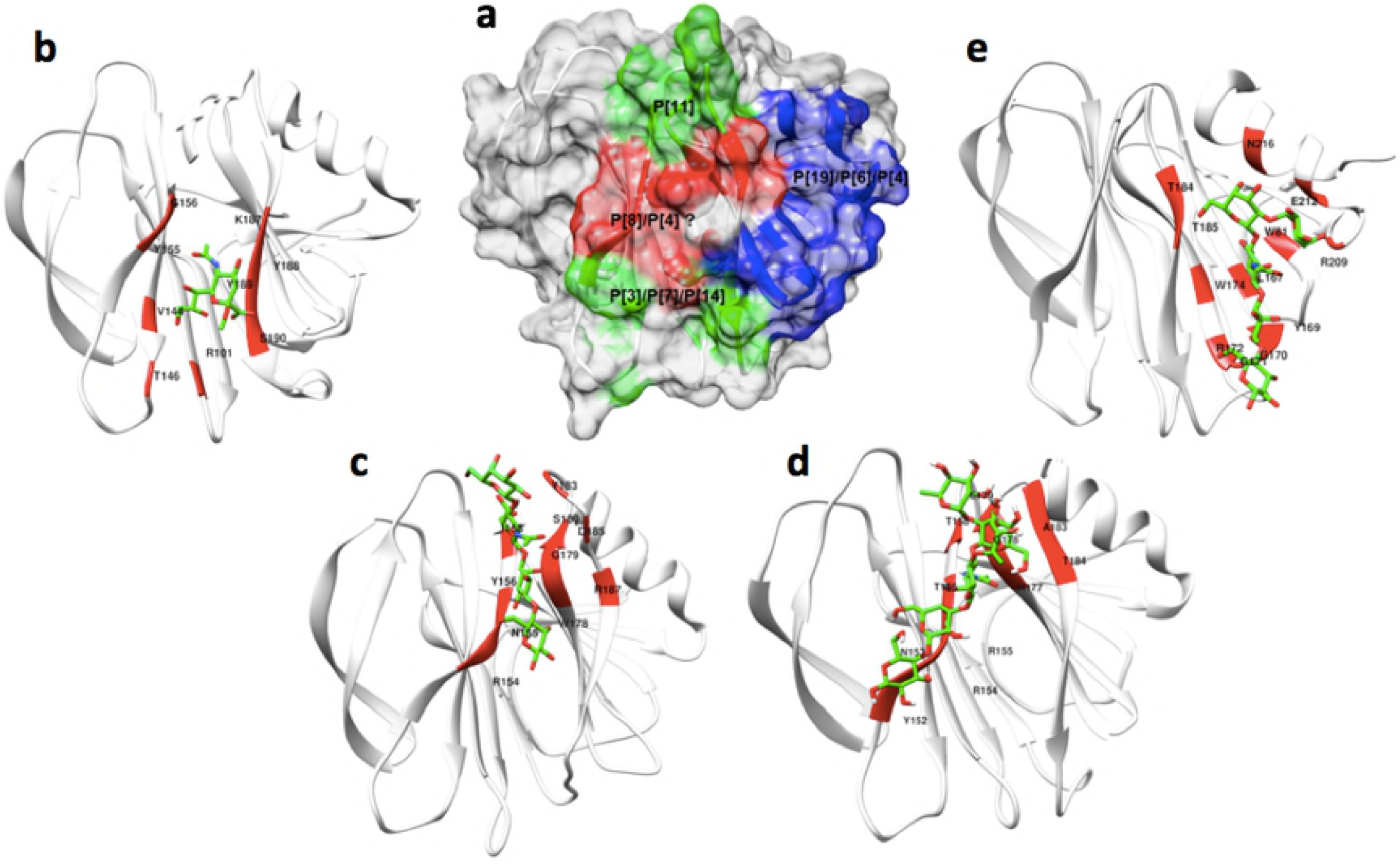
The binding interface of different genotypes of VP8* in recognizing their corresponding glycans. (a) Surface-rendered structure of XXX depicting a summary of the distinct glycan receptor binding sites identified to date. The red-colored region indicates the **ββ** binding motif that P[3]/P[7] use in binding sialic acid, P[14] in binding type A HBGAs, P[11] in binding type 1 HBGAs, and P[8] in recognizing Lewis type HBGAs. The blue-colored region indicates the **βα** binding pocket that the P[19], P[6] and P[4] RVs use in recognizing H-type 1 HBGAs. (b) Detail of the sialic acid binding interface (red) of P[3] [PDB ID:1KQR]. (c) Detail of the LNT binding site of P[11] [PDB ID: 4YFZ]. (d) Detail of the LNDFH I binding interface of P[8]. (e) Detail of the LNFP I binding site of P[4] [PDB ID: 5VX5].

## Discussion

Recently, the importance of HBGA as cell attachment factors has been demonstrated for multiple different human RV genotypes, however, the molecular level understanding of the glycan binding interactions of P[8] RVs to HBGAs remains incompletely understood and under continued investigation. Huang originally reported that the P[8] genotype used Lewis and H type 1 HBGAs as ligands for RV attachment [19], which have been observed by others [43]. However, other groups contradicted these results, reporting that human P[8] strain does not recognize Lewis b and H type 1 antigens [44]. In an attempt resolve these contradictory results, we have used saturation transfer difference and HSQC-based NMR titration experiments to study the RV VP8* -ligand interactions. Our results confirmed that human P[8] VP8* can bind both Lewis b and H type 1 HBGAs. As an alternative to X-Ray crystallography, we used NMR-driven HADDOCK docking to generate data-driven models of the complexes of the P[6] and P[8] VP8* domains with their respective glycan ligands, which is widely used to characterize protein-ligand interactions when crystals of the complex of interest are unavailable, which is commonly the case for weak binding systems [38–40].

Given the fact that the multiple genotypes in P[II] genogroup exhibit variable host ranges among animal species and different human populations, further clarification of the molecular basis for ligand-controlled host ranges of the major human RVs is important for development of improved vaccines in the future. Structural-based sequence alignment of P[19], P[6], P[4] and P[8] showed that they have high levels of amino acid conservation. From the recent crystallization studies, it was found that P[19], P[6], and P[4] employed the βα binding domain to accommodate H type 1 ligand LNFP I [35,36], which is in agreement with our NMR-driven HADDOCK results. Previous homology modeling results suggested that the addition of Lewis fucose via α1,4-linkage to the O4 atom of the GlcNAc residue could cause a binding clash if P[8] used the same binding interface as P[4] [36]. Our structural studies presented here resolve this perceived conflict, showing that P[8] accommodates the Le^b^-containing ligands by shifting to a new binding site, namely the ββ binding domain. Therefore, unlike P[4], P[6] and P[19], P[8] uses the ββ binding domain to accommodate Le^b^ tetra-saccharide and LNDFH I, which appears to reflect an evolutionary adaptation driven by selective pressure to accommodate the addition of the Lewis epitope to the HBGAs. Furthermore, based on our STD NMR results, we observed that P[4] can also recognize the Le^b^-containing LNDFH I ligand (refer to supplementary figure). Although we do not yet have conformational experimental structural data, we hypothesize that P[4] may also use the ββ binding domain to accommodate the Le^b^-containing LNDFH I ligand. Structural studies to test this hypothesis are currently underway in our laboratories.

Our data also provide clues about the evolution of P[II] RVs with a hypothesis under strong selection by type 1 HBGAs. While most P[I] RVs infect animals, the majority of P[II] RVs infect humans. P[II] RV was hypothesized to have originated from a P[I] RV with a possible animal host origin and then gained the ability to infect humans by adapting to the polymorphic HBGAs in humans [45]. P[19] RVs are commonly found to infect animals (porcine) but rarely humans. P[19] RVs exhibit binding specificities to mucin core 2 and type 1 HBGA precursors, indicating that P[19] genotype may be positioned at an early evolutionary stage starting from a common P[I] ancestor. Recently, the apo crystal structure of porcine P[6] strain (z84) and human P[6] strain (RV3) have been reported, both demonstrating a galectin-like fold as reported for P[19] [36,46]. P[6] RVs are commonly found to infect animals (porcine), neonates and young infants [47.48]. Expression and modification of the precursor moieties in the neonate gut is developmentally regulated, and unbranched type 1 precursor glycans are the most abundant type in the neonate gut [31,49]. It was found from our NMR titration experiments that P[6] had a much higher affinity to the tetra-saccharide LNT than the penta-saccharide LNFP I, supporting the previous findings that P[6] had a preference to less mature type 1 HBGA precursors. Furthermore, P[6] RVs have an age-specific host range and exhibit cross-species transmission between humans and animals, indicating P[6] may represent an RV evolutionary intermediate that adapted to limited glycan residues shared between porcine and humans. Compared to P[6], P[4] and P[8] are genetically closely related and their infections are more commonly detected. The binding affinity of P[8] to the hexa-saccharide LNDFH I is much tighter than to the Le^b^ tetra-saccharide based on the NMR titration data, supporting the view that P[8] and P[4] are more distantly evolved from their P[I] ancestor and have adapted to recognize more mature type 1 HBGAs. The fact that P[4] and P[8] are commonly found in humans but rarely in animals would be consistent with their ability to bind additional residues in more complex HBGAs such as is found in the incorporation of Lewis epitopes [25], which are widely distributed in humans but not commonly expressed in animals, such as mice [50].

The elucidation of HBGA-correlated susceptibility to infection with specific RV P-genotypes, mainly associated with secretor status and Lewis phenotype, is important to understand RV disease burden and epidemiology. HBGAs are fucose-containing carbohydrates and act as receptors for various pathogenic microorganisms, such as *Helicobacter pylori* [51], Norovirus [52], and Rotavirus. Six types of precursor oligosaccharide chains have been characterized, and type 1 can be found in the epithelium of the respiratory, gastrointestinal, genitourinary tracts and in exocrine secretions [24]. The expression of polymorphic human HBGAs are genetically regulated by specific glycosyltransferase, and the resultant HBGA products for the ABO, H and Lewis families are distributed among the world’s population with their frequencies varied from one ethnicity to another [53]. For example, the addition of α-1,2 fucose is catalyzed by the α-1,2 fucosyltransferase that encoded by the FUT2 gene, which is the major determinant of secretor antigens. H-positive phenotypes who contain secretor antigens occurs in about 80% of European and North American populations [54]. The Lewis-positive individuals who contain an active α-1,3 fucosyltransferase (FUT3) enzyme represent about 90% of the general population, but the frequencies is much lower in Africa [55]. The discovery that P[4], P[6], and P[8] human RVs recognize the secretor epitopes of human HBGAs appears to correlated with the predominance of these genotypes in causing the vast majority (>95%) of human infections worldwide. Epidemiological and biochemical studies suggest that non-secretors and Le^-^ individuals may be resistant to P[4] and P[8] infections [27,28,56,57]. Non-secretors are also possible to be susceptible to infection with prevalent human RVs as other epidemiological study demonstrated no significant correlation was found between secretor status and the susceptibility to P[6] RV infections [26]. In the current study, we demonstrated that P[6] recognized the tetra-saccharide LNT that contains the type 1 HBGA sequences without secretor and Lewis epitopes, and the penta-saccharide LNFP I that contains H epitopes but no Lewis epitopes. Compared with human P[4] and P[8] that recognize Lewis epitope, P[6] HRVs bind H type 1 only and have a restricted geographic prevalence and are common in the African countries [58,59]. In these countries, the higher prevalence of P[6] HRVs could be due to the significantly higher rate of Le negative phenotype among the population than in other geographic locations [12,28].

Even though all of the solved VP8*structures share the same conserved galectin-like fold, they exhibit significant glycan recognition variability in a genotype, i.e. evolutionary, dependent manner. While differential immune responses, presence of other co-receptors and variable host factors may all account for the relative distribution of RV genotypes and variable vaccine efficacy in different populations [56], achieving a more complete understanding of the specificity of glycan recognition specificity of VP8*domains may help understand the limited effectiveness of the two current RV vaccines in certain populations and provide clues for the formulation of more effective vaccines. For example, based on relative effectiveness of the two current RV vaccines, Rotarix and RotaTeq, in the developing versus developed countries and their different vaccine designs, a role of the P type (VP4/VP8*) in host immune protection against RVs has been emphasized, which has led to a hypothesis on the lack of cross-protection of the P[8] based Rotarix and RotaTeq against other RV P types that are more commonly seen in the developing countries than in the developed countries could be a major reason of the low effectiveness of both vaccines observed in many developing countries (refer to our recent review article, Jiang et al 2017), although other factors, such as malnutrition, intestinal microbiota, human maternal milk of children living in many developing countries may also play roles. Thus, based our emerging understanding of RV diversity, strain-specific host ranges, and RV evolution, an approach that includes other P types in RV vaccines, such as the P[6] and P[11] RVs that are more commonly seen in the developing countries, may improve the global effectiveness of the current vaccines.

## Materials and methods

### Expression and purification of VP8* proteins in *Escherichia coli*

The VP8* core fragments (amino acids 64 to 223) of the human RV P[8] (BM13851) and P[6] (BM11596) with an N-terminal glutathione S-transferase (GST) tag was overexpressed in *Escherichia coli* BL21 (DE3) cells as previously described [19]. Cells were grown in 1L Luria broth (LB) medium supplemented with 100 μg ml^-1^ ampicillin at 310 K. When the OD600 reached 0.8, 0.5 mM isopropyl-β-D-thiogalactopyranoside was added to the medium to induce protein expression. The cell pellet was harvested within 12 h after induction and re-suspended in the phosphate-buffered saline (PBS) buffer (140 mM NaCl, 2.7 mM KCl, 10 mM Na_2_HPO_4_, 1.8 mM KH_2_PO_4_, pH 7.3). The cells were lysed by French press (Thermo Fisher Scientific, Waltham, MA), and the supernatant of the bacterial lysate was loaded to a disposable column (Qiagen, Hilden, German) pre-packed with glutathione agarose (Thermo Fisher Scientific). The GST fusion protein of interest was eluted with elution buffer (10 mM reduced glutathione, 50 mM Tris-HCl, pH 8.0). The GST tag of the VP8* protein was removed using the thrombin (Thermo Fisher Scientific) after dialysis into the buffer (20 mM Tris-HCl, 50 mM NaCl, pH 8.0). The VP8* protein without GST tag was further purified by passing the mixture through size exclusion chromatography using a Superdex 200 Hiload (GE Life Science) column. The purified protein was concentrated with an Amicon Ultra-10 (Millipore, Billerica, MA) for future NMR study. The ^15^N or ^15^N, ^13^C-Labled P[8] and P[6] proteins was made using the ^15^N or ^15^N,^13^C-Labeled minimal growth medium.

### Ligand chemical shift assignments

Chemical shift assignments of Le^b^ tetra-saccharide, lacto-N-tetraose (LNT), and lacto-N-difucohexaose I (LNDFHI) were completed by collecting and analyzing the following NMR spectra at 20°: 2D ^1^H-^13^C heteronuclear single quantum correlation spectroscopy (HSQC), 2D ^1^H-^13^C heteronuclear multiple-bond correlation spectroscopy (HMBC), ^1^H-^1^H correlation spectroscopy (COSY), ^1^H-^1^H total correlation spectroscopy (TOCSY), ^1^H-^1^H rotating-frame nuclear Overhauser effect spectroscopy (ROESY), and ^1^H-^13^C HSQC-TOCSY. All the glycans were prepared in PBS buffer (pH 7.3). The chemical shifts of lacto-N-fucopentaose I (LNFPI) were previously assigned [25].

### Saturation transfer difference (STD) NMR experiments

All STD NMR spectra were acquired in Shigemi Tubes (Shigemi, USA) on 600-MHz Bruker Avance III and 850-MHz Bruker Avance II NMR spectrometers equipped with conventional 5-mm HCN probes at 283 K with pulse sequence STDDIFFESGP.3. The NMR samples for P[8] VP8* were prepared as 46 μM P[8] VP8* with 2.3 mM LNDFHI (1:50 protein: ligand ratio), and 33 μM P[8] VP8* with 1.86 mM Le^b^ tetra-saccharide (1:56 protein: ligand ratio). The NMR samples for P[6] VP* were prepared as 46 μM P[6] with 4.6 mM LNT (1:100 protein: ligand ratio), and 46 μM P[6] with 2.3 mM LNT (1:50 protein: ligand ratio). A control sample was prepared as 40 μM GST with 2.0 mM LNDFHI (1:50 protein: ligand ratio). The STD NMR spectrum of glutathione S-transferase (GST) mixed with LNDFH I was collected as a negative control and it did not show significant signal intensities as expected, validating the specificity of our results. The protein resonances were saturated with a cascade of Gaussian-shaped pulses with a duration of 50 ms, with a total saturation time of 2 s and 4 s. The on resonance was chosen to selectively saturate only the proteins for each sample. The off-resonance saturation was applied to 50 ppm, and a total of 256 scans were acquired. A spin-lock filter with 20 ms duration was applied to suppress the broad protein resonance signals, and residual water signal was suppressed using excitation sculpting with gradients. The STD amplification factors were determined by the equation (I_STD_/I_O_) × molar ratio, where I_0_ are the intensities of the signals in the reference spectrum, and I_STD_ are intensities that obtained by subtracting the on-resonance spectrum from the off-resonance spectrum. The normalized STD amplification percentage for epitope mapping was determined by dividing by the largest STD amplification factor for each ligand.

### Protein backbone chemical shift assignment

The ~0.6 mM double-labeled ^15^N, ^13^C-P[6] and P[8] VP8* sample were put into 5-mm Shigemi NMR tubes and spectra were collected at 298 K on 600-MHz Bruker Avance III and 850-MHz Bruker Avance II NMR spectrometers equipped with conventional 5-mm HCN probes. Backbone assignments were made based on the following three-dimensional spectra: HNCACB, CBCA(CO)NH, HNCO, HN(CA)CO, HNCA, and HN(CO)CA. Spectra were processed with the software NMRPipe [60] and visualized with the software SPARKY [61] (https://www.cgl.ucsf.edu/home/sparky/). The backbone ^1^H, ^13^C, and ^15^N resonance assignments were first made using the PINE server [62], then manually confirmed through SPARKY.

### NMR titration experiments

Chemical shift perturbations were followed by 2D ^1^H-^15^N HSQC for P[8] VP8* and P[6] VP8* upon titration of each glycan ligand. NMR data were collected with samples in PBS buffer in 5-mm Shigemi NMR tubes on 600-MHz Bruker Avance III or 850-MHz Bruker Avance II NMR spectrometers equipped with conventional 5-mm HCN probes. Samples of 0.20 mM ^15^N-labeled P[8] VP8* was titrated with LNDFHI at 0, 0.020, 0.040, 0.80, 1.00, 1.20, 1.40, 1.60, 2.00, 2.80, 3.60, 4.40, 5.00 mM. Samples of 0.31 mM ^15^N-labeled P[8] VP8* was titrated with Le^b^ tetra-saccharide at 0, 0.031, 0.15, 0.31, 0.62, 1.24, 2.49, 4.35, 5.60, 6.84, 7.78, 8.71, 9.33 mM. Samples of 0.24 mM ^15^N-labeled P[6] VP8* was titrated with LNFP I at 0, 0.024, 0.12, 0.24, 0.48, 0.96, 1.93, 3.37, 4.34, 5.30, 6.03, 6.75, 7.23, 8.44 mM. Samples of 0.33 mM ^15^N-labeled P[6] VP8* was titrated with LNT at 0, 0.033, 0.17, 0.33, 0.69, 1.34, 2.67, 4.68, 6.01, 7.35, 8.35 mM. Spectra were process with NMRPipe software [60] and analyzed with SPARKY [61]. The chemical shift changes upon titration was determined by the following formula [63]: 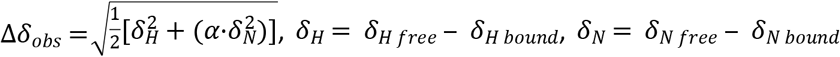. Where *δ_H free_* and *δ_H bound_* are the backbone amide hydrogen chemical shifts in the absence and presence of ligand, *δ_N free_* and *δ_H bound_* are the amide nitrogen chemical shifts in the free and bound state. The value of Euclidean weighting correction factor α was set to 0.14 [64]. Standard deviation σ of the Euclidean chemical shift change was calculated and threshold value was chosen from 1.5σ to 2.5σ for the VP8*-HBGA by balancing the specificity and sensitivity. Amino acids with chemical shifts greater than the threshold value were used to calculate the dissociation constants. Dissociation constants were obtained by fitting the following equation: 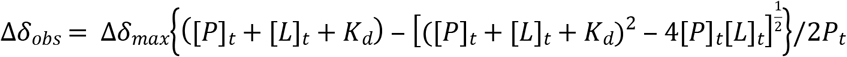. Where [*P*]*_t_* and [*L*]*_t_* represent the total concentrations of protein and ligand, Δ*δ_obs_* is the change in the observed chemical shift from the free state, Δ*δ_max_* is the maximum chemical shift change on saturation, *K_d_* represents the dissociation constant.

### Homology modeling and HADDOCK docking

A homology model of the P[8] (BM13851) VP8* protein was built based on the X-ray crystal structure of the Wa strain of RV (PDB: 2DWR), and a homology model of the P[6] (BM11596) VP8* protein was built based on the X-ray crystal structure of the z84 strain of RV (PDB: 5YMU). The model building was finished by SWISS-MODEL automated protein structure homology-modeling server [65]. Docking simulation were calculated with HADDOCK2.2 [40,66]. Ambiguous interactions restraints were defined for active residues in protein identified to be involved in the interactions using NMR titration, and with high solvent accessibility. Solvent accessibility of the proteins were performed by freeSASA [67], and amino acids with solvent accessibility of both main chain and side chain less than average values are filtered out. STD data were used to refine the ambiguous interaction restraints for ligands, and trNOESY data were used to constrain the conformation of the bound ligand. trNOESY were performed at 1:20 protein/ligand ratio and at a mixing time of 100ms, 150ms, and 200 ms. Longitudinal cross-relaxation rates were obtained by averaging the normalized volume at the three mixing times. From them, intramolecular ligand proton-proton distances were obtained by using the isolated spin pair approximation, and taking the distance of GlcNAc H1-H2 as reference. Each docking calculation generated 1000/200/200 models for the rigid body docking, semi flexible simulated annealing, and explicit solvent refinement. To optimize protein-ligand docking, the RMSD cutoff for clustering was set to 2.0, the Evdw 1 and Eelec 3 were set to 1.0 and 0.1 respectively, and the initial temperatures for second and third TAD cooling step were set to 500 K and 300 K respectively. The model with the lowest/best HADDOCK score (a linear combination of various energies and buried surface area) in the best cluster was picked for further analyze and visualization using Chimera [68] and LigPlot+ [69]. Sequence alignment was finished by Clustal Omega [70].

